# Multimodal imaging brain markers in early adolescence are linked with a physically active lifestyle

**DOI:** 10.1101/2020.09.03.269571

**Authors:** Piergiorgio Salvan, Thomas Wassenaar, Catherine Wheatley, Nicholas Beale, Michiel Cottaar, Daniel Papp, Matteo Bastiani, Sean Fitzgibbon, Euguene Duff, Jesper Andersson, Anderson M. Winkler, Gwenaëlle Douaud, Thomas E. Nichols, Stephen Smith, Helen Dawes, Heidi Johansen-Berg

**Author notes:** Corresponding author: Piergiorgio Salvan, Wellcome Centre for Integrative Neuroimaging, Nuffield Department of Clinical Neurosciences, University of Oxford, Oxford OX3 9DU, United Kingdom.

## Abstract

The World Health Organization (WHO) promotes physical exercise and a healthy lifestyle as means to improve youth development. However, relationships between physical lifestyle and brain development are not fully understood. Here, we asked whether a brain – physical latent mode of covariation underpins the relationship between physical activity, fitness, and physical health measures with multimodal neuroimaging markers. In 50 12-year old school pupils (26 females), we acquired multimodal whole-brain MRI, characterizing brain structure, microstructure, function, myelin content, and blood perfusion. We also acquired physical variables measuring objective fitness levels, 7-days physical activity, body-mass index, heart rate, and blood pressure. Using canonical correlation analysis we unravel a latent mode of brain – physical covariation, independent of demographics, school, or socioeconomic status. We show that MRI metrics with greater involvement in this mode also showed spatially extended patterns across the brain. Specifically, global patterns of greater grey matter perfusion, volume, cortical surface area, greater white matter extra-neurite density, and resting state networks activity, covaried positively with measures reflecting a physically active phenotype (high fit, low sedentary individuals). Showing that a physically active lifestyle is linked with systems-level brain MRI metrics, these results suggest widespread associations relating to several biological processes. These results support the notion of close brain-body relationships and underline the importance of investigating modifiable lifestyle factors not only for physical health but also for brain health early in adolescence.

**Significance statement:** An active lifestyle is key for healthy development. In this work, we answer the following question: How do brain neuroimaging markers relate with young adolescents’ level of physical activity, fitness, and physical health? Combining advanced whole-brain multimodal MRI metrics with computational approaches, we show a robust relationship between physically active lifestyles and spatially extended, multimodal brain imaging derived phenotypes. Suggesting a wider effect on brain neuroimaging metrics than previously thought, this work underlies the importance of studying physical lifestyle, as well as other brain – body relationships in an effort to foster brain health at this crucial stage in development.

## Introduction

The World Health Organisation (WHO) encourages early positive lifestyle choices aimed to improve both physical and mental health (World Health Organization 2010). Physical activity is a powerful and rapid means to improve fitness and physical health throughout the life-span (Cotman 2002; Charles H. Hillman, Erickson, and Kramer 2008). During adolescence, however, levels of physical activity decline (Guthold et al. 2020).

Public health guidelines recommend that school-aged children engage in 60 minutes of moderate-to-vigorous physical activity daily (Piercy and Troiano 2018), yet globally only around 22% of boys and 15% of girls achieve that (Guthold et al. 2020). In addition to its importance to physical health, there is growing evidence that a physically active lifestyle during childhood is associated with improved mental and cognitive health through adulthood (Promotion and US Department of Health & Human Services Office of Disease Prevention and Health Promotion 2000). While there is limited available evidence in adolescents, similar patterns have been reported (Lubans et al. 2016).

A body of work has studied the relationship between single physical measures of activity, fitness, or body-mass, and separate MRI metrics of brain structure, microstructure, or function, showing focal neural correlates (for reviews see (Valkenborghs et al. 2019; Donnelly et al. 2016)). However, it is unlikely that a single physical measure fully captures active lifestyles, or that a single MRI metric fully quantifies the condition of the brain. Rather, lifestyles are better characterized by a range of physical measures and the state of the brain is better quantified by combinations of metrics.

Multimodal MRI can probe different aspects of brain structure and function. While each metric provides an indirect probe of the underlying biology, in combination they provide insights into a range of biological processes (Tardif et al. 2016). Further, these measures can be acquired simultaneously across the whole brain. Many previous brain imaging studies of physical activity and fitness have focused on the hippocampus, where changes in non-invasive imaging measures of tissue volume or perfusion have been argued to relate to processes of neurogenesis and angiogenesis triggered by exercise (van Praag et al. 1999; Pereira et al. 2007; Chaddock et al. 2011; Thomas et al. 2012). However, in addition to such focal changes, more global biological processes might also be triggered by exercise (Tardif et al. 2016). It remains unknown whether whole-brain patterns of multimodal brain metrics are related to cardiorespiratory fitness, physical activity, and physical health.

Physical activity influences physical health and contributes to physical fitness but both activity and fitness may be considered part of an underlying, latent factor. In order to characterize a phenotype of physical lifestyle, measuring whole-day physical activity levels during a normal school week is therefore at least as important as assessing gold-standard measures of cardiorespiratory fitness, such as VO2max measured on an incremental step-test on a cycle ergometer.

In this study, in 50 12-year old pupils, we acquired multimodal whole-brain MRI metrics to measure resting state networks (RSNs), grey matter (GM) volume and perfusion, cortical surface (area and thickness), white-matter (WM) microstructure, and myelin content (R1 and R2*), resulting in a total of 18 different metrics). These metrics are combined into multimodal whole-brain phenotypes whose variation across individuals can be interrogated. We also acquired a rich set of variables depicting physical lifestyle, measuring cardiorespiratory fitness (VO2max and workload), objective physical activity (7-days actigraphy, measuring total week time of brief bursts and long-lasting physical activity) and reported (questionnaire item), and physical health (resting heart rate, blood pressure, and body-mass index) (**Fig. 1**). We hypothesised that across pupils, inter-subject differences in brain phenotypes covaried with differences in physical lifestyle, independent of sex, socioeconomic status, age, pubertal level, and school. A single holistic multivariate analysis allowed us to identify a latent mode of covariation between brain and physical phenotypes, representing a pattern of active physical lifestyle features that significantly covaries with spatially extended patterns of brain metrics.

**Figure 1.**
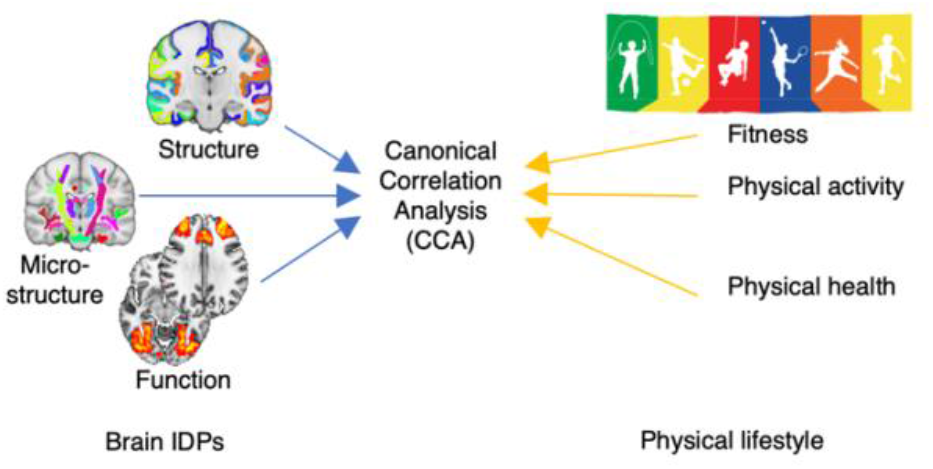
Summary of statistical analysis. In order to test the individual covariation between brain IDPs and physical measures of fitness, physical activity and physical health, we aimed to identify one single mode of covariation using canonical correlation analysis (CCA), while taking into account the hierarchical structure represented by schools.

## Materials and methods

### Participants

Year 7 pupils from a subset of 10 UK schools participating in the Fit to Study project, were invited to take part in a brain imaging substudy (Wassenaar et al. 2019). After taking consent and assent in accordance with the University of Oxford ethical guidelines (CUREC reference number: R51313/RE001), 61 pupils were recruited to the brain imaging substudy. Participants attended a testing session at the University of Oxford during which brain imaging, cognitive, and behavioral data were collected.

Only 50 pupils (median age: 12 years; 26 females (52%)) (**Table 1**), had high-quality, complete multimodal MRI data. All statistical analyses were carried out on this sample of 50 pupils sampled from 10 schools (**Table 2**).

**Table 1.**
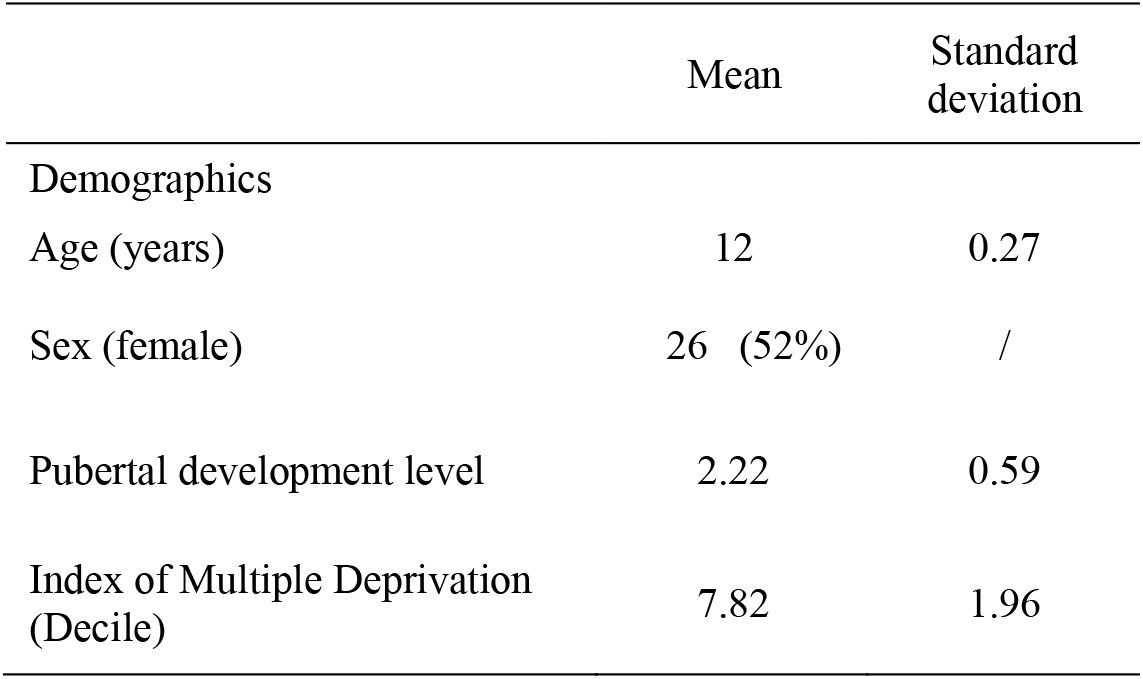
Demographics and socioeconomic status.

|  | Mean | Standard deviation |
| --- | --- | --- |
| Demographics |  |  |
| Age (years) | 12 | 0.27 |
| Sex (female) | 26 (52%) | / |
| Pubertal development level | 2.22 | 0.59 |
| Index of Multiple Deprivation (Decile) | 7.82 | 1.96 |

**Table 2.** Sampling frequency by school.

| School | Number of pupils | % |
| --- | --- | --- |
| A | 1 | 2 |
| B | 2 | 4 |
| C | 10 | 20 |
| D | 6 | 12 |
| E | 8 | 16 |
| F | 1 | 2 |
| G | 1 | 2 |
| H | 1 | 2 |
| I | 13 | 26 |
| L | 7 | 14 |

### Behavioral testing

#### Cardiorespiratory fitness

Objective measures of *cardiorespiratory fitness* were acquired through an incremental step-test on a cycle ergometer (Lode Excalibur Sport, Groningen, The Netherlands). We then extracted values for maximal oxygen consumption per kilogram (VO2/kg max) (ml/min/kg), and work load maximum (Watts) as primary measures of interest.

#### Physical activity

Objective *physical activity* was assessed over five weekdays and 2 weekend days using the Axivity AX3 wrist-worn accelerometer (Open Lab, Newcastle University, UK) (Ladha et al. 2013). We therefore chose to define a valid wear day as 12 consecutive hours from 08.00 to to capture travel to and from school and after-school sports and activities. To account for later weekend waking times, we accepted any consecutive ten-hour period between 08.00 and 20.00 on Saturdays and Sundays, and standardised total activity to 12 hours. We then aimed to capture both *brief bursts* and *long-lasting* activity. We converted raw accelerometer data into activity ‘counts’ per 60-second epoch and also per 1-second epoch to characterize sustained bouts of activity and also shorter bursts of movement. Axivity’s Open Movement GUI software calculated whether each 60-second epoch was spent in sedentary, light, moderate or vigorous activity by applying established adolescent cut-points (Phillips, Parfitt, and Rowlands 2013). The software identified non-wear time as periods of at least 30 consecutive minutes of zero activity counts. We used a bespoke programme, designed to handle large volumes of data, to apply the same cut-points to each 1-second epoch. Participants who had at least 3 valid weekdays and one valid weekend day were included in the analysis (Troiano et al. 2014). For both *brief bursts* and *long-lasting* physical activity, participants’ total minutes of sedentary, moderate and vigorous activity per day were calculated.

#### Physical health

*Physical health* was assessed on the day of testing at rest (prior cardiorespiratory testing) by measuring heart rate; systolic and diastolic blood pressure. When compared to publicly available age-matched normative values (Flynn JT et al. 2018), blood pressure (5th to 95th percentiles) was found within healthy values (normative values for 12 years old pupils: *systolic*: 102-131 with average of 113; *diastolic*: 61-83 with average of 75; study sample, *systolic*: median = 106, 5th-95th percentiles: 89-123; *diastolic*: median = 73, 5th-95th percentiles: 58-85).

#### Negative behaviors not considered in the analysis

As part of the study we also obtained ethics to ask pupils information about negative behaviors such as smoking, drinking alcohol, or drug use. However, none of the pupils reported having used any of these substances.

### MR Imaging

#### MRI acquisition parameters

All Magnetic Resonance Imaging (MRI)-scans were carried out during summer 2017 at the Oxford Centre for Functional MRI of the Brain (FMRIB) using a 3T Siemens Magnetom Prisma (Erlangen, Germany) scanner with a 32-channel head coil.

The MRI protocol included:

1. T1 weighted (T1w) three-dimensional rapid gradient echo sequence (3D MPRAGE): repetition time (TR) = 1900 ms; echo time (TE) = 3.97 ms; flip angle = 8°; field-of-view (FOV) = 192 mm; voxel size: 1× 1 × 1 mm. Sequence duration: 5 min 31 sec.
2. Resting-state functional MRI (rs-fMRI): multi-band echo-planar imaging (EPI) sequence; TR = 933ms; TE = 33.40 ms; FOV = 192 mm; 72 slices; voxel size: 2 × 2 × 2 mm; multi-band acceleration factor = 6. Sequence duration: 10 min 10 sec. For each scan, 644 volumes were acquired. Participants were asked to look at a fixation cross, blink normally, try not to fall asleep and not to think about anything in particular. A field map was also acquired to correct for inhomogeneity distortions. Sequence duration: 1 min 34 sec.
3. Diffusion-weighted MRI (DW-MRI): multi-shell, multi-band EPI sequence; b-values = 0, 1250, 2500 s/mm2, with respectively 11, 60, 60 diffusion-weighted directions; TR = 2483 ms; TE = 78.20 ms; FOV = 214 mm; voxel size: 1.75 × 1.75 × 1.75 mm; multi-band acceleration factor = 4. Sequence duration: 5 min 40 sec. In addition, 4 b = 0 s/mm_2_ images were acquired with reversed phase encoding, for the purpose of EPI distortion correction. Sequence duration: 32 sec.
4. Quantitative FLASH-MRI (Weiskopf et al. 2013): two 3D multi-echo FLASH datasets, one predominantly proton-density weighted (PDw, flip angle = 6 deg), and one predominantly T1w (flip angle = 21 deg); FOV = 256 mm; voxel size: 1× 1 × 1 mm; TR = 25 ms; first TE = 2.34 ms; eight equally space echoes, echo spacing = 2.3; GRAPPA acceleration factor = 2 in both phase-encoded directions, with 40 reference lines in each direction. Duration for each FLASH sequence: 5 min 11 sec. Two single-echo, low-resolution (4 mm isotropic) FLASH scans were acquired before each high-resolution scan; identical FOV; TR = 4 ms; TE = 2 ms; one was acquired receiving on the 32-channel receive head coil, the other receiving on the body coil. To correct for the effect of RF inhomogeneities, the local RF field was mapped using a 2D DAM method with a FLASH readout.
5. Pseudo-continuous arterial spin labeling (pCASL) with background pre-saturation (Okell et al. 2013): six imaging blocks, each with different post-labeling delays: 0.25, 0.5, 0.75, 1, 1.25 and 1.5 s. Arterial blood was magnetically tagged using a labeling duration of 1.4 s. Other imaging parameters were: single shot EPI; TR = 4100 ms; TE = 14 ms; FOV = 220 mm; voxel size: 3.4 × 3.4 × 4.5 mm. Sequence duration: 5 min 34 sec.

### MRI preprocessing

MRI data was processed primarily using FSL software (Jenkinson et al. 2012) and FreeSurfer (Dale, Fischl, and Sereno 1999).

#### Gradient distortion correction

Gradient distortion correction (GDC) was applied within image analysis pipelines using tools developed by FreeSurfer and HCP groups (https://github.com/Washington-University/Pipelines), using the Siemens scanner specific table of gradient nonlinearities.

#### Structural

Brain extraction was performed in native space after GDC unwarping using FSL BET (Smith 2002). Tissue-type segmentation was estimated based on FSL FAST (Zhang, Brady, and Smith 2001), providing hard segmentation as well as partial-volume images for each tissue type. This tool was also used to provide a fully bias-field-corrected version of brain extracted structural brain images. Subcortical structures were modelled using FSL FIRST (Patenaude et al. 2011).

#### Cortical surface reconstruction

Subject-specific cortical surface reconstruction and cortical parcellation were estimated based on the GDC, brain extracted T1 image, using the command *recon-all* from FreeSurfer (Dale, Fischl, and Sereno 1999).

#### Registration

Rigid registrations between multimodal MRI native spaces were estimated through FSL FLIRT with boundary-based cost function (Jenkinson et al. 2002; Greve and Fischl 2009). Nonlinear warps to MNI152 standard-space T1 template were estimated through FSL FNIRT. This set of nonlinear warps is then carried over to all MRI modalities such as in the case of resting state functional MRI (rs-fMRI).

#### EPI distortion correction

B0 fieldmap processing was estimated through FSL Topup (Andersson, Skare, and Ashburner 2003) based on AP-PA image pairs from DWI-MRI protocol.

#### Functional

rs-fMRI data was preprocessed using a custom pipeline previously validated on developmental datasets (Baxter et al. 2019). rs-fMRI data was corrected for inter- and intra-volume subject head motion, EPI distortions (Andersson and Sotiropoulos 2015); highpass temporal filtering and GDC unwarping was also applied. Registration to structural was improved by an extra rigid registration step aided by a single-band EPI image. Structured artefacts were removed by FSL ICA+FIX processing (Beckmann and Smith 2004; Salimi-Khorshidi et al. 2014; Griffanti et al. 2014). The FSL FIX classifier was specifically trained for this data and provided the following scores in leave-one subject-out accuracy: true positive ratio (TPR) = 98.8%; true negative ratio (TNR) = 95.3%; weighted ratio ((3*TPR+TNR)/4) = 97.9%. FSL MELODIC was then used to estimate 50 group-average independent components (ICs). We then calculated median absolute (ridge) partial correlation (with a regularization value of 0.1) and amplitude for each of the 25 ICs identified as RSNs. *Diffusion.* DWI-MRI data was first corrected for eddy currents, EPI distortions, inter- and intra-volume subject head motion, with outlier-slice replacement, using FSL Eddy (Andersson and Sotiropoulos 2015). GDP unwarping was then applied (Miller et al. 2016). Diffusion Tensor Imaging (DTI) fitting was carried out with FSL DTIFIT using a kurtosis model (Behrens et al. 2007). Neurite Orientation Dispersion and Density Imaging (NODDI) modelling was estimated using FSL cuDIMOT (https://users.fmrib.ox.ac.uk/~moisesf/cudimot/DesignModel.html) based on the Bingham-NODDI model (Tariq et al. 2016). In order to resolve crossing-fibres configurations, multi-shell voxel-wise diffusion was modelled using FSL BedpostX (Jbabdi et al. 2012). Probabilistic tractography was then carried out with FSL ProbtrackX (Behrens et al. 2007) and 29 major WM bundles were reconstructed as implemented in FSL AutoPtx (de Groot et al. 2013) _31_.

#### Myelin and iron maps

Quantitative MRI data was processed to produce the quantitative maps of myelination (1/T1) and iron level (1/T2*), using the Voxel-Based Quantification (VBQ) toolbox (Callaghan et al. 2014) in Statistical Parametric Mapping (SPM) (http://www.fil.ion.ucl.ac.uk/spm/). Although R1 (1/T1, longitudinal relaxation rate) and R2* (1/T2*, effective transverse relaxation rate) are not *direct* quantitative maps of myelination or iron (as other biological factors can also affect them), these quantitative maps have high degree of sensitivity to myelination and iron (Weiskopf et al. 2013; Lutti et al. 2014; Callaghan et al. 2014).

#### Perfusion

Perfusion images were processed using FSL BASIL (Chappell et al. 2009). Images were first corrected with fieldmap and GDC unwarping; then, in order to obtain maps of cerebral blood flow and arrival time in absolute units, a calibration step was implemented based on cerebrospinal fluid values.

### Image derived phenotypes (IDPs)

Each MRI parameter was summarized in a series of IDPs: anatomy-specific average values that span three sets of regions of interest. For cortical and subcortical regions, we used the Desikan-Killiany Atlas (84 parcels, cortical and subcortical) from the individual FreeSurfer parcellation (Fischl et al. 2002). This parcellation was then warped into each (relevant) modality in order not to interpolate MRI-map values. For the white matter, we used the 29 white-matter bundles from the AutoPtx reconstruction; first averaged at group-level; optimally thresholded; and then warped back to native spaces. For functional activity, 25 group-level RNSs were identified.

A total of 859 IDPs were then fed into statistical analysis.

### Cognitive testing and reported mental health and general health measures

All measures acquired during the testing are reported in detail in (Wassenaar et al. 2019).

### Cognitive skills

Here we considered a summary measure for 3 tasks of interest: the relational memory task (correct valid answers (%)) (Chaddock et al. 2010); task switching (switch cost (ms)) (C. H. Hillman et al. 2014); and object-location task (identification errors 8s-delay) (Pertzov et al. 2012), (**Table 3**).

**Table 3.** Descriptives of cognitive skills, reported mental health, and reported general health.

|  | Mean | Standard deviation |
| --- | --- | --- |
| Cognition |  |  |
| Associative task (correct valid answers (%)) | 71 | 11 |
| Task switching (switch cost (ms)) | 548 | 333 |
| Object location task (Identification errors 8s-delay) | 11 | 4 |
| Mental health (SDQ questionnaire) |  |  |
| Prosocial Scale | 8 | 1.4 |
| Hyperactivity Scale | 4 | 2.5 |
| Conduct Scale | 2 | 1.5 |
| Peer Scale | 2 | 1.9 |
| Emotional Scale | 3 | 2.3 |
| General Health (HSBC questionnaire) |  |  |
| Life satisfaction | 8 | 1.6 |
| Self-Rated Health | 4 | 0.9 |
| Multiple health complaints | 13 | 4.8 |

### Mental health

Mental health was assessed with the Strengths and Difficulties Questionnaire (SDQ) (Goodman 1997).

### Questionnaire on general health

From the Health behavior in school-age children (HSBC) questionnaire *(World Health Organization. Regional Office for Europe 2016)* we used the positive health items (self-rated health, life satisfaction, multiple health complaints) in order to measure reported general health.

### Experimental design and statistical analyses

This is a cross-sectional study with a sample size of N = 50 subjects. Due to the limited sample size compared to the number of variables of interest, we strove to reduce input data and nuisance variables dimensions as much as possible. All statistical analyses were carried out in MATLAB 2018.

### Confounds

Prior to all statistical analyses, a series of relevant confounds was chosen: age; sex; pubertal developmental level (assessed through the *Pubertal Development Rating Scale* (Petersen et al. 1988), a self-report measure of physical development for youths under the age of 16); socioeconomic status (assessed through the UK Index of Multiple Deprivation); and head size/scaling factor (computed through FSL SIENAX).

On these nuisance variables we perform a dimensionality reduction through means of principal component analysis (PCA) (Nz = 2) accounting for 60% of total variance. These confounds were then regressed out of all IDPs and behavioral variables and the residuals normalised.

### Dimensionality reduction of IDPs and physical variables

In order to avoid an overdetermined, rank deficient CCA solution, and to limit the chances of overfitting, a dimensionality reduction set was performed to both IDPs and physical variables. Using the same approach previously applied in (Smith et al. 2015), IDPs were reduced into 10 PCAs (Nx = 10; variance explained = 53%), whilst physical variables were reduced into 5 PCAs (Ny = 5; variance explained = 79%).

### Canonical correlation analysis

We sought to characterize a mode of brain – physical covariation across pupils: a data-driven latent factor linking a linear combination of neuroimaging metrics (**Table 4**) with a linear combination of physical measures (**Table 5**). To this end we used canonical correlation analysis (CCA), an approach that has successfully been applied in recent studies and that, compared to pairwise association testing, has shown greater sensitivity for complex biological processes and greater explained variance (Miller et al. 2016; Smith et al. 2015). CCA is a symmetric, cross-decomposition method that characterizes covariation modes between a pair of two-dimensional datasets. This is achieved by finding two sets of free parameters (or *canonical coefficients*, i.e. one set of coefficient vectors per set of brain metrics and one set of coefficient vectors per set of physical metrics) that maximize the correlation of the projections of the two datasets into the identified latent space (or *canonical variates* or subject *scores*). In other words, the variation in mode strength between subjects is maximally correlated. Here, this was computed using MATLAB ‘canoncorr’ function.

**Table 4.**
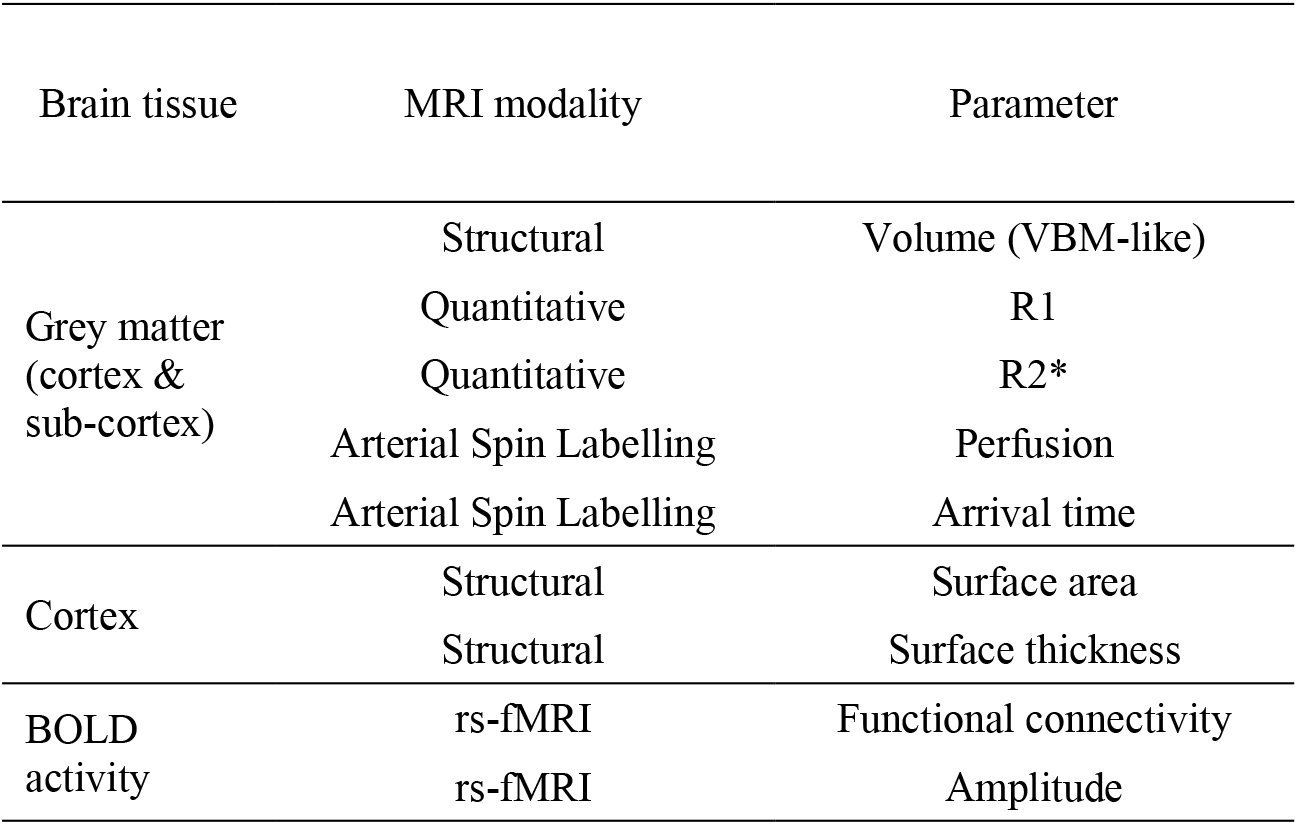
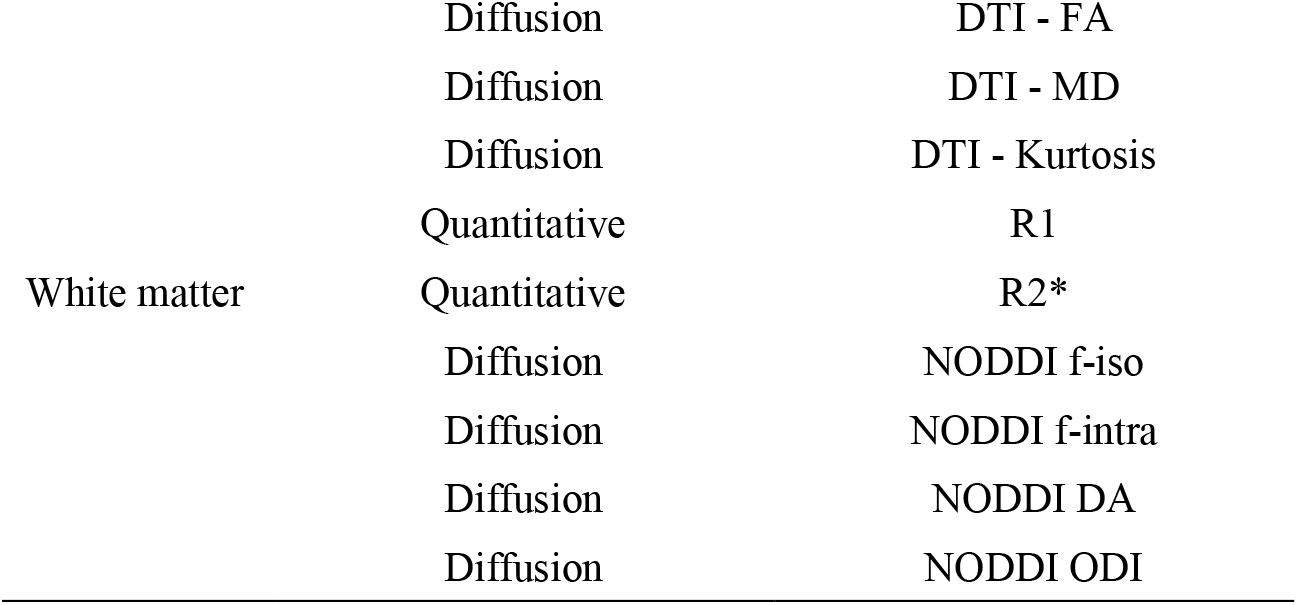
List of MRI modalities and MRI metrics used to define IDPs of brain function and structure. 5 MRI sequences were used to quantify 18 different MRI metrics. Specific sets of ROIs were then used for each MRI metric in order to extract whole-brain MM IDPs quantifying brain structure, microstructure, function, myelin content, and blood perfusion.

**Table 5.**
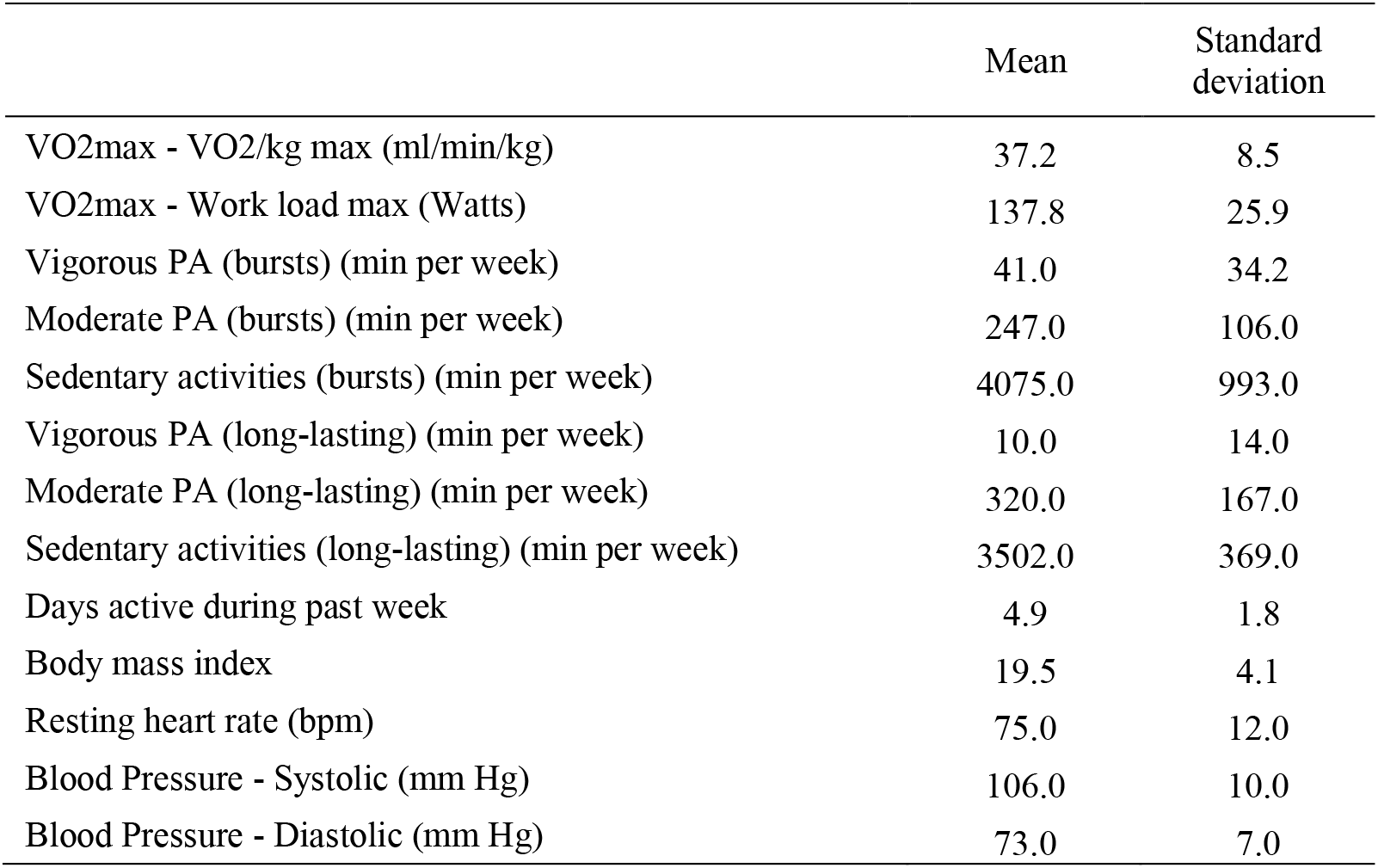
Descriptives of physical variables. 13 measures of physical activity, fitness, and physical health were considered in testing the relationship with brain IDPs. Here we report mean and standard deviation (prior of correcting for demographics and socioeconomic status).

|  | Mean | Standard deviation |
| --- | --- | --- |
| VO2max - VO2/kg max (ml/min/kg) | 37.2 | 8.5 |
| VO2max - Work load max (Watts) | 137.8 | 25.9 |
| Vigorous PA (bursts) (min per week) | 41.0 | 34.2 |
| Moderate PA (bursts) (min per week) | 247.0 | 106.0 |
| Sedentary activities (bursts) (min per week) | 4075.0 | 993.0 |
| Vigorous PA (long-lasting) (min per week) | 10.0 | 14.0 |
| Moderate PA (long-lasting) (min per week) | 320.0 | 167.0 |
| Sedentary activities (long-lasting) (min per week) | 3502.0 | 369.0 |
| Days active during past week | 4.9 | 1.8 |
| Body mass index | 19.5 | 4.1 |
| Resting heart rate (bpm) | 75.0 | 12.0 |
| Blood Pressure - Systolic (mm Hg) | 106.0 | 10.0 |
| Blood Pressure - Diastolic (mm Hg) | 73.0 | 7.0 |

### Unbiased statistical inference through block-aware permutation testing

Deconfounding, as required to ensure that the CCA is not driven by nuisance factors, induces a dependency among the rows of the data submitted to CCA. While this dependency is weak and diminishes with increasing sample size, it represents a violation of the exchangeability assumption required by permutation, that can inflate permutation significance. To account for this deconfounding-induced dependency that violates exchangeability, we use a method that, without changing the canonical correlations, reduces the data from N observations to N-Nz observations that are exchangeable, and thus, can be subjected to a permutation test (Winkler et al. 2020; Theil 1965). We randomly chose 1,000 sets of Nz rows for removal, conducting 1,000 permutations for each set.

Permutations were performed among subjects within school, respecting dependencies given by the hierarchical structure of the data (Winkler et al. 2015). For each of the 1,000 repetitions, a *p-*value was computed based on this null distribution for the first CCA mode. Across repetitions, a distribution of statistical significance values was built and the final statistical significance level was computed as its average value. The results of this analysis are shown in **Fig. 2b**.

**Figure 2.**
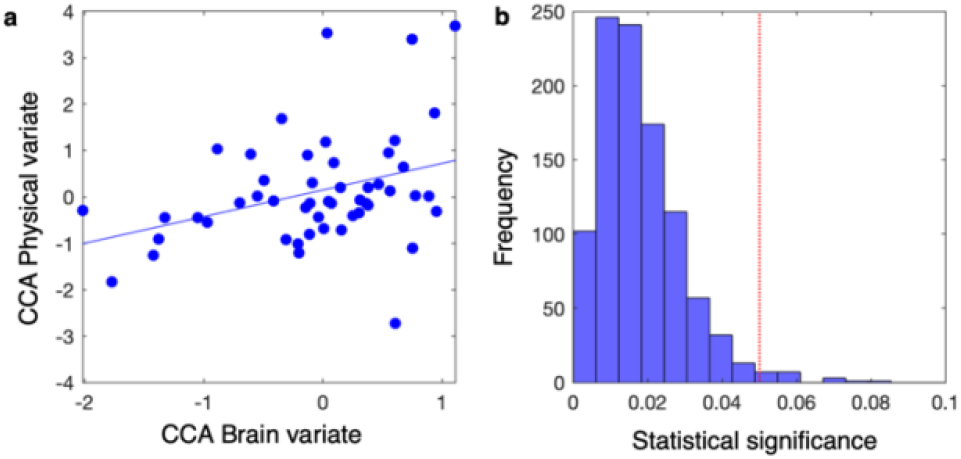
Mode of brain – physical covariation across pupils. The results from canonical correlation analysis (CCA) highlight one significant mode of brain – physical covariation across pupils. **a**) Showing scatter plot of cross-validated canonical variates between brain IDP scores and physical scores, each dot represents a pupil (cross-validated CCA: rho = 0.34). Statistical significance of CCA was assessed 1,000 times, each time comparing the real value against 1,000 block-aware permutations taking into account school structure. **b**) Showing distribution of statistical significance values. The final significance value was assessed as the average of this distribution (p-value = 0.0130). Red dashed line represents cut-off of statistical significance of alpha = 0.05.

### Unbiased estimation of effect size through leave-one school-out cross-validation

In order to derive an unbiased estimate of the CCA correlation strength that took into account the hierarchical structure in the data, we implemented a leave-one school-out cross-validation (CV) approach. In all but one school, we performed all the above steps (except permutation testing), learning all the coefficients of the linear transformations. On the left-out school, we then applied those transformations and predicted left-out pupils' scores in the CCA mode. We repeated this procedure for all folds (here schools). CV performance was then quantified as the Pearson’s rho correlation coefficient and mean squared error (MSE) calculated between predicted brain and physical canonical covariates (or predicted canonical variates or CCA subject *scores*). The results of this analysis are shown in **Fig. 2a**.

### Characterization of brain and physical phenotypes

We then aimed to characterize the CCA phenotypes: the set of brain measures and the set of physical measures symmetrically linked by the CCA covariance mode. To do this (formally, to characterize the CCA crossed *loadings*), we follow the procedure described in (Smith et al. 2015). On the whole sample, CCA Brain loadings were calculated as the pairwise Pearson’s *partial* correlation between CCA Physical variate (or subject *scores*) and the original datasets of brain IDPs, while controlling for the full set of nuisance variables (CCA Brain loadings = partialcorrelation(brain IDPs, CCA Physical variate, nuisance variables). The results of this process are shown in **Fig. 4**, **5**, **6**. CCA Physical loadings were calculated with the following the same process (CCA Physical loadings = partialcorrelation(physical variables, CCA Brain variate, nuisance variables). The results of this process (only for structural IDPs) are shown in **Fig. 3**. CCA loadings are therefore bounded between 1 and −1.

**Figure 3.**
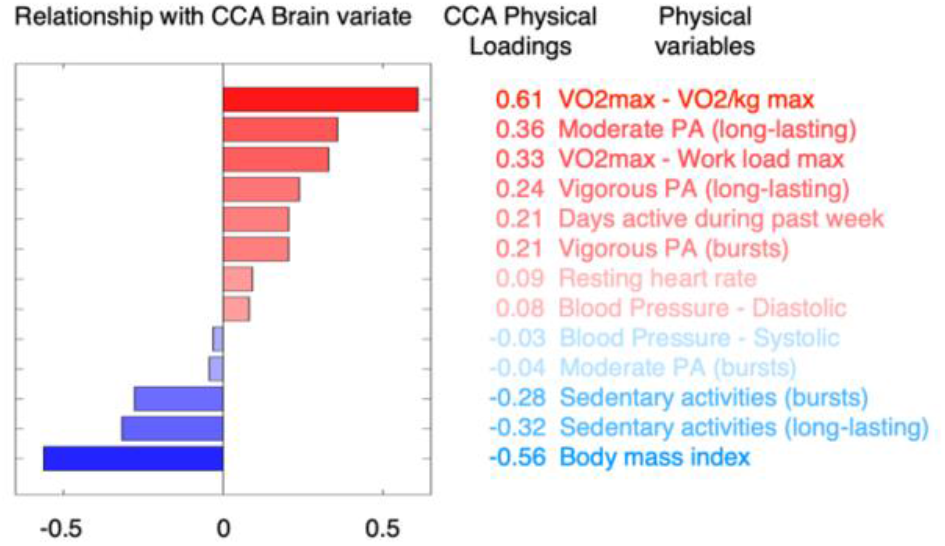
Physical phenotype linked to the brain – physical mode of covariation. Bar plot representing the coefficient structure of physical phenotype (formally CCA Physical loadings). Each coefficient represents the relationship between each physical metric and subjects’ brain IDP scores (or CCA Brain variate). Bar plot and variable ranking are matched and color-coded in red/blue in accordance to a positive/negative relationship with the mode of covariation (the magnitude of involvement is further represented through transparency).

**Figure 4.**
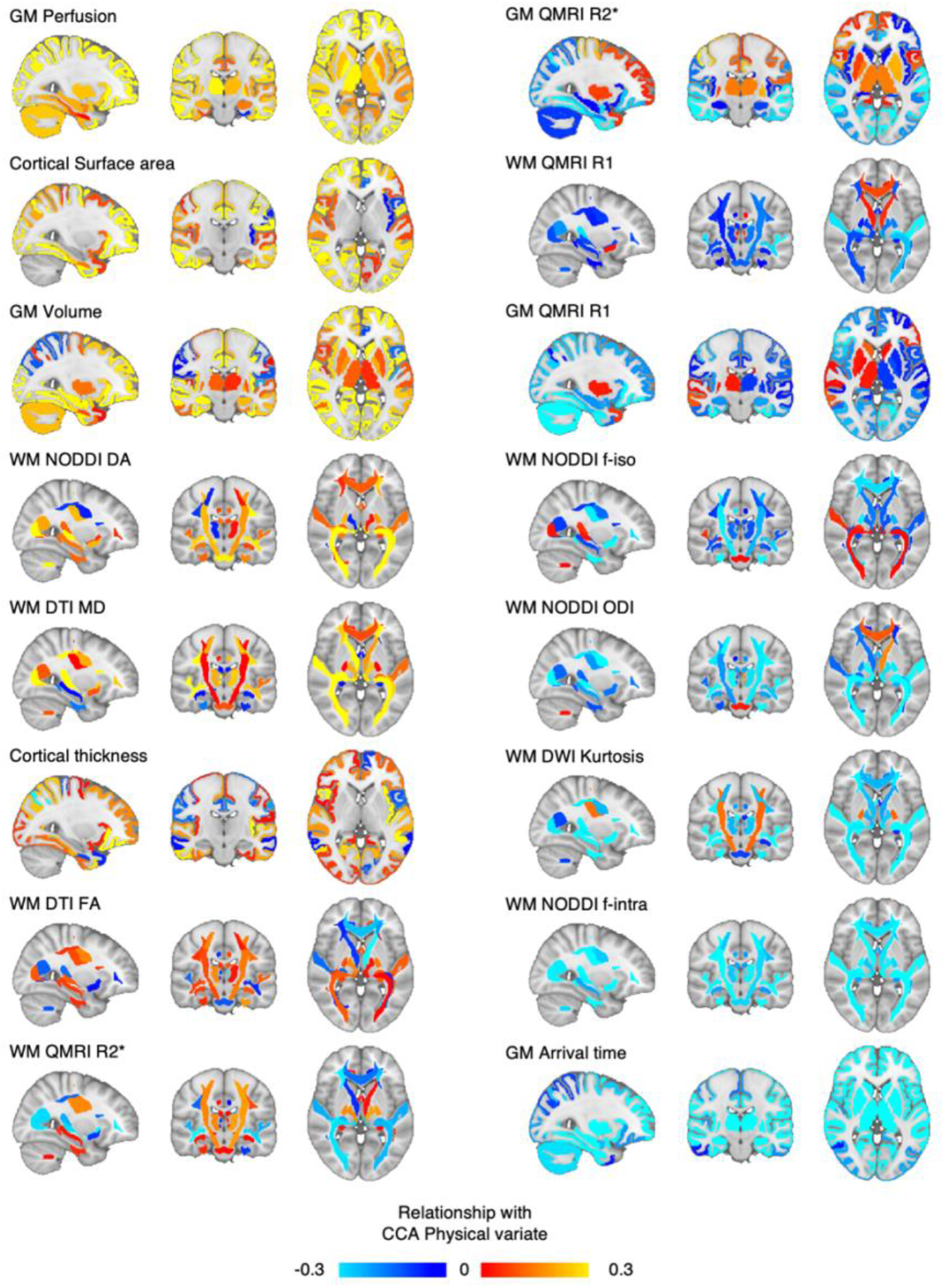
Structural IDPs and their relationship with the identified phenotype of physically active lifestyle. Showing for each MRI metric and for each structural ROI, thus for all structural IDPs, the relationship with the identified phenotype of active lifestyle. Hot colors represent a positive relationship with the physical phenotype; cold colors represent a negative relationship. Structural maps are ranked from top to bottom (left column to right) in accordance to *average* CCA W Brain.

**Figure 5.**
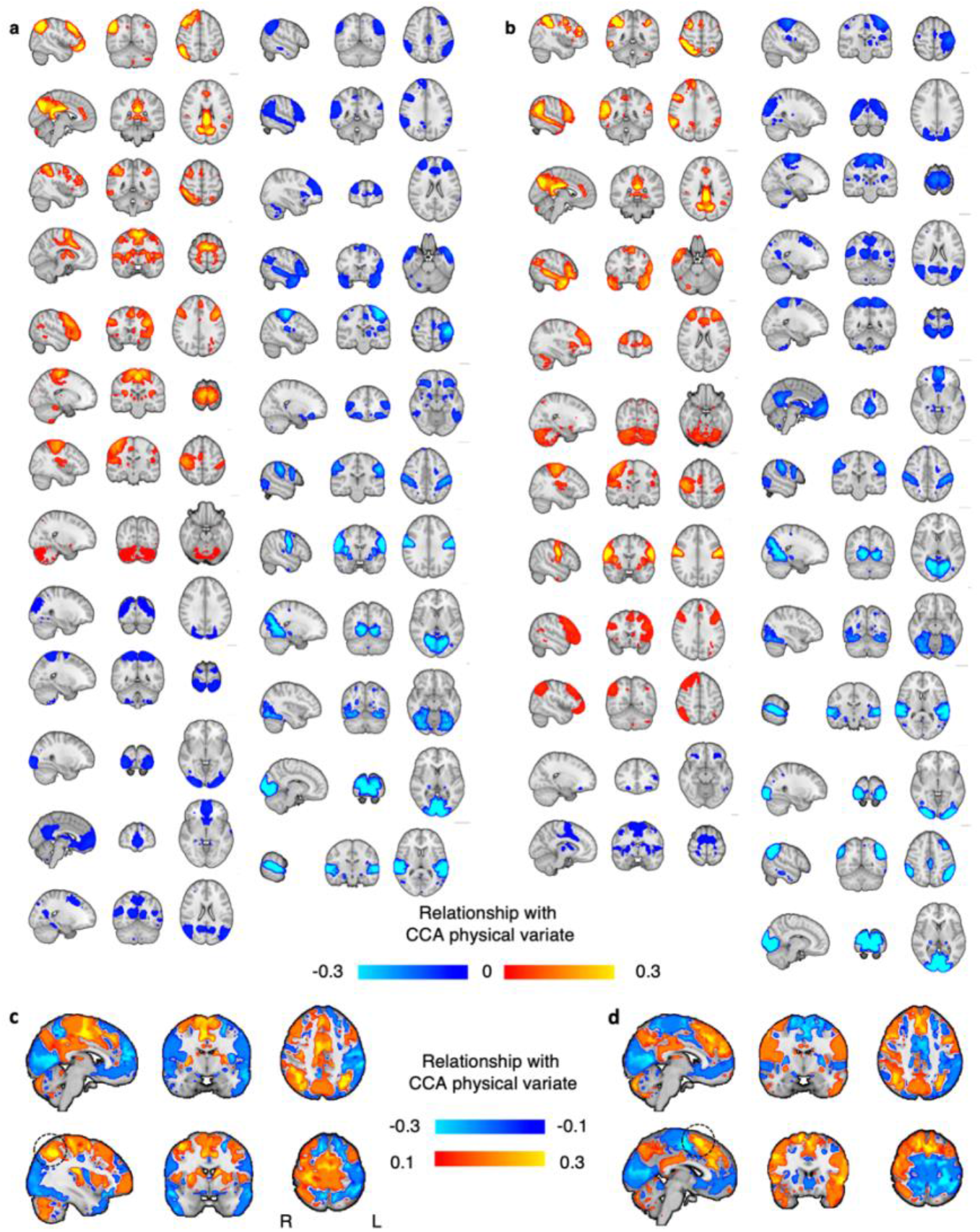
Functional IDPs and their relationship with the identified phenotype of physically active lifestyle. Showing for each resting state brain network (RSN), and for both metrics functional connectivity **a**) and amplitude **b**), the relationship with the identified phenotype of active lifestyle. Hot colors represent a positive relationship with the physical phenotype; cold colors represent a negative relationship. To aid interpretation, for each RSN, its CCA Brain loading was multiplied by the group RSN map. RSN are here ranked from top-to-bottom in accordance to their CCA Brain loadings. We then concatenated all RSNs maps in a 4D file and computed the mean and standard deviation across RSNs, separately for both functional connectivity and amplitude. **c**) Normalised mean of CCA Brain loadings for RSNs functional connectivity. **d**) Normalised mean of CCA Brain loadings for RSNs amplitude. For **c**) and **d**), the top row shows the same brain coordinates, whilst the bottom row shows the respective peak of greater CCA Brain loadings. Dashed circle in **c**) and **d**) represents the peak value.

**Figure 6.**
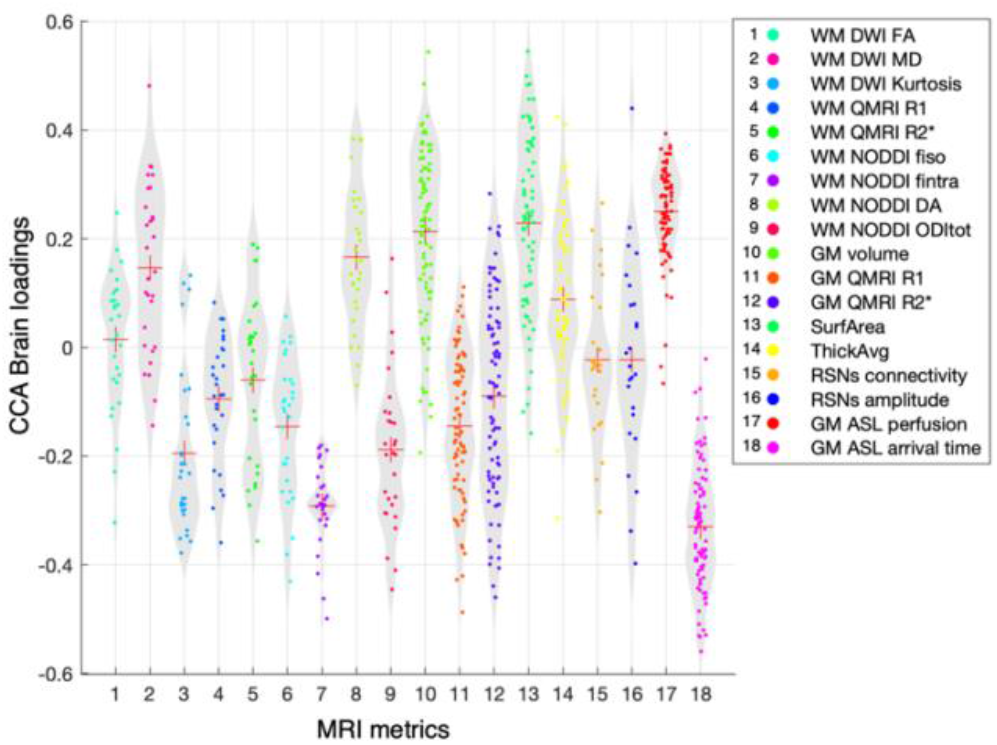
Relationship with physical lifestyle phenotype for all brain IDPs. Showing the CCA Brain loadings for all 859 brain IDPs divided into each MRI metric. Each dot represents one single IDP.

For functional measures, each IDP is represented by a whole-brain RSN. To aid interpretation, for each RSN, its CCA Brain loading was multiplied by the group RSN map. The results of this process are shown in **Fig. 5a, b**. Then, in order to derive a summary representation, we concatenated all RSNs maps in a 4D file and computed normalised mean across RSNs, separately for both functional connectivity and amplitude. The results of this process are shown in **Fig. 5c, d**.

We then aimed to characterize the average involvement for each type of MRI value. Across IDPs of a MRI metric, we computed the average across CCA Brain loadings. This provided a ranked list of MRI parameters representing the average relationship of each MRI metric with pupils’ physical scores. The results of this process are shown in **Fig. 7**.

**Figure 7.**
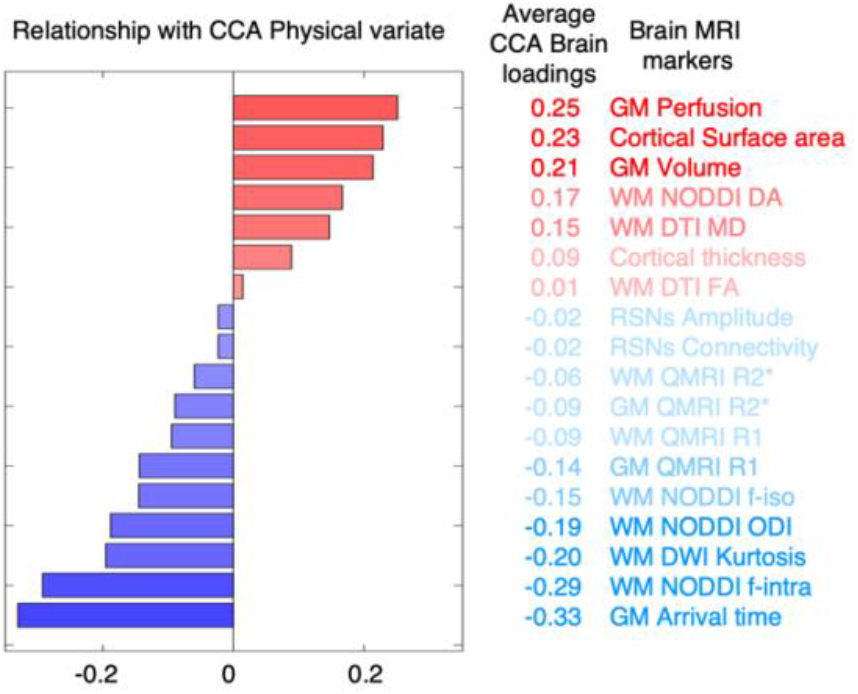
Brain phenotype linked to the brain – physical mode of covariation. Bar plot representing the *average* CCA Brain loadings. Each coefficient represents the relationship between each MRI metric (average across ROIs) and pupils’ physical lifestyle scores. Bar plot and variable ranking are matched and color-coded in red/blue in accordance to a positive/negative relationship with the mode of covariation (the magnitude of involvement is further represented through transparency).

### Joint-inferences with univariate measures of cognitive skills, mental health, and general health

The relationships between the identified CCA mode and the multiple variables measuring the domains of cognitive skills, mental health, and general health, were carried out (separately for each domain) with a general linear model with non-parametric combination (NPC) implemented in FSL PALM (Winkler et al. 2016). NPC works by combining test statistics or p-values of separate (even if not independent) analyses into a single, joint statistic, the significance of which is assessed through synchronized permutations for each of the separate tests. Here we asked whether the CCA mode was associated with any sub-measure within a domain, and the NPC was tested via Tippett statistic with 1,000 block-aware permutations, while adjusting for nuisance variables in reduced space.

## Results

### Brain – physical mode of covariation across pupils

Using CCA, we tested the hypothesis that across pupils, inter-subject differences in multimodal whole-brain IDPs covaried significantly with differences in physical lifestyle variables, independent of nuisance variables. We found one significant mode of brain – physical covariation across pupils, linking differences in brain IDPs with individual differences in physical lifestyle (**Fig. 2a**; CCA: rho = 0.34, MSE = 1.38, using leave-one school-out CV; **Fig. 2b**; p-value = 0.0130, significance assessed on 1,000 repetitions, each with 1,000 block-aware permutations; results remained the same if adjusted for the full set of nuisance variables). This mode represents a pattern of brain IDPs that covaries with a pattern of physical variables. We next interrogated this physical phenotype and brain phenotype separately, to determine the patterns that underlie this mode.

### Physical phenotype of covariation

For each physical variable we calculated the loadings of the physical phenotype relating to the CCA mode (or ‘CCA Physical loadings’) representing the relationship between each physical variable and the CCA brain variate (or *subjects’ brain scores*) (**Fig. 3**). We found that pupils who scored higher in the brain – physical mode of covariation were those with higher cardiovascular fitness; those with lower body-mass index; those who spent more time doing long-lasting (both moderate and vigorous) physical activity during a normal school week and spent less time being sedentary.

### Brain phenotypes of covariation

In order to interpret brain phenotypes of physical covariation, we calculated the canonical loadings for each IDP of brain structure, microstructure, and function. These loadings (or ‘CCA Brain loadings’) represent the relationship between each brain IDP and the CCA Physical variate (or *subjects’ physical scores*) (to explore the spatial patterns of *all* structural and functional IDPs, see respectively **Fig. 4** and **Fig. 5a, b**; for violin plot of CCA Brain loadings for all IDPs, see **Fig. 6**).

We observed that some MRI metrics presented a global and homogeneous involvement in the mode of covariation across ROIs. In order to quantify this tendency, for each MRI metric, we computed the *average* CCA Brain loadings across all ROIs (**Fig. 7**). We found that the strongest CCA Brain loadings were found for GM perfusion (and arrival time) as well as cortical surface area, GM volume and a number of WM diffusion metrics. MRI metrics with the greatest average CCA Brain loadings tended to be characterized by spatially extended and homogeneous involvement across the whole-brain. Together, these results show that pupils with greater physical scores were those who also showed global patterns of higher blood perfusion (and lower arrival time, i.e., faster perfusion) in the GM, greater GM volume, greater cortical surface area, greater neurite dispersion anisotropy across WM tracts, as well as greater extra-neurite fraction (equivalent to lower *intra*-neurite fraction), and lower neurite orientation dispersion.

We also observed that although the *average* CCA Brain loadings for RSNs functional connectivity and BOLD amplitude was close to zero, there was great variance across RSNs (**Fig. 5**). Because RSNs are not binary masks but are instead characterized by spatial distributions, in order to summarise their pattern of involvement in the mode of covariation, for each voxel we computed the normalised mean CCA Brain loadings across RSNs (**Fig. 5c, d**). The resulting maps for functional connectivity (**Fig. 5c**) and amplitude (**Fig. 5d**) showed both similarities and differences in their patterns of involvement in the mode of covariation. In Fig. 5.c, RSN functional connectivity shows greater positive involvement bilaterally in the parietal cortices, supplementary motor cortex, putamen, and right primary motor cortex; while it shows greater negative involvement broadly in the occipital cortices. The peak of positive involvement was localised in the right parietal cortex, while the negative in the occipital cortex. In Fig. 5d, RSN BOLD amplitude shows greater positive involvement in the anterior cingulate gyrus (dACC), superior frontal gyrus, parietal cortices, right inferior frontal gyrus; while it shows greater negative involvement broadly in the occipital cortices, and left primary somatosensory cortex. The peak of positive involvement was localised in the dACC, while the negative in the occipital cortex. These maps show a common pattern of greater positive involvement bilaterally in the parietal cortices, and a common pattern of negative involvement in the occipital cortices.

### Relationship with measures of cognition, mental health, and general health

We then tested the hypothesis that the identified CCA mode of brain – physical covariation (averaging between brain and physical CCA subjects scores), was significantly associated with measures of 1) cognitive skills, 2) mental health, and 3) general reported health. Testing a NPC joint-inference for each domain, we found no statistically significant association (NPC Tippett p-value respectively for cognitive skills: p-value = 0.3280; mental health: p-value = 0.7940; general health: p-value = 0.1280).

Of interest for future studies, we report that one of the three HBSC questionnaire items in general reported health (*self-rated health*) was positively associated with the mode of brain – physical covariation if not corrected for multiple comparisons across the three items tested (Adjusted R2 = 0.16, p-value = 0.0430).

Importantly, when not using block-aware permutations and thus not taking into account the hierarchical structure represented by schools, the association between the mode of brain – physical covariation and the domain of general health was found to be significant. This highlights the importance of considering schools' effect (or any other hierarchical structure) when testing associations across a population. It also emphasizes how robust the results from the brain – physical covariation are to nuisance of no interest and to hierarchical structure in the data.

## Discussion

In this work we show that, in 12-year-old pupils, physical activity, fitness, and physical health are linked with global patterns of brain structure, microstructure, and function. In this relationship, whole-brain, homogeneous patterns of multimodal brain phenotypes are linked with a specific, latent pattern of physical measures that capture a physically active lifestyle (high fit, high active, low sedentary individuals). This finding hints at the involvement of multiple underlying biological processes and suggests that physical health and aerobic exercise might have a wider effect on brain processes than previously thought.

We applied a holistic approach to provide novel insight into the importance of different aspects of a physically active lifestyle in relation to brain structure and function. While high cardiovascular fitness and physical activity are positively linked with the identified brain phenotypes, sedentary activity and body-mass index are negatively related. Furthermore, we showed that long-lasting physical activity, either moderate or vigorous, is more important to this relationship than brief bursts of activity, suggesting that regular moderate-to-vigorous physical activity might be a better driver to promote brain changes. Taken together, these findings situate pupils along a latent axis according to their physical phenotype: pupils with high cardiorespiratory fitness and performance and with high weekly levels of physical activity, g, contrast with pupils spending most time in sedentary or low-energy behaviors. The novelty of this work is the finding of multimodal global brain phenotypes linked with a physically active lifestyle. Although prior work has studied the relationship between single measures of brain structure or function and, separately, physical activity or fitness (Valkenborghs et al. 2019), our approach allowed us to identify latent patterns of multimodal brain IDPs characterized by the involvement of multiple brain regions in the covariation with physical scores. Specifically, greater physical scores were linked with spatially extended patterns of greater blood perfusion and faster arrival time in the GM, greater GM volume and larger cortical surface area, and in the WM with lower intra-neurite density and kurtosis. This result shows that high fitness and physical activity are associated with more *global* patterns of brain structure than previously thought. Further work is needed to better understand multimodal, spatially extended phenotypes of brain structure (Groves et al. 2012; Douaud et al. 2014). Indeed it remains unknown how spatially extended brain patterns relate to individual differences in cognition, their level of heritability, as well as to what extent they are susceptible to plasticity. Although it is not possible to infer the presence of a specific biological process or cellular component solely on the basis of MRI measures (Zatorre, Fields, and Johansen-Berg 2012), these results suggest that high fitness and regular physical activity might have a more *widespread* impact on brain structure than previously thought.

Previous literature has explicitly focused on studying the effects of aerobic exercise on the hippocampus (Cotman 2002; Falkai and Schmitt 2009; Chaddock-Heyman et al. 2016; Thomas et al. 2016). Cardiorespiratory fitness is indeed known to promote hippocampal neurogenesis and angiogenesis that, in turn, determines macro-scale changes that are also visible via non-invasive neuroimaging (van Praag et al. 1999). Here, we extend the current knowledge beyond a uniquely hippocampal pattern, highlighting the global nature of greater volume and faster perfusion across the whole brain GM. In other words, variation in hippocampal structure alone does not underlie the brain – physical relationship characterized here. Rather we observed homogeneous loadings across GM areas. The strongest contributions to our brain phenotype came from perfusion measures. These robust associations found with perfusion metrics are in line with a body of literature showing positive effects of physically active lifestyle on vascular health (Vaynman, Ying, and Gomez-Pinilla 2004; Promotion and US Department of Health & Human Services Office of Disease Prevention and Health Promotion 2000), as well as animal studies linking physical exercise to angiogenesis (Kleim et al. 1996; Rhyu et al. 2010).

Further key insights derive from spatially extended patterns of WM covariation. Although the myelin sensitive metrics (Quantitative-MRI R1 and R2*) in the current study made little contribution to the mode of variation, higher scores on the physical phenotype were associated with lower intra-neurite density and kurtosis, and to a lesser extent with lower neurite orientation dispersion and with greater dispersion anisotropy. It is relevant that the DW-MRI protocol employed in this study would be sensitive to diffusion properties within large glial cells, such as astrocytes and oligodendrocytes. Our gradient strength provides sensitivity to length-scales of roughly 4-6 μm, with the body-size of astrocyte and oligodendrocytes being respectively in the order 20 μm (Oberheim et al. 2009) and of 14 μm (Bakiri et al. 2011), much larger than the average myelinated axon diameter (<1 μm) (Liewald et al. 2014). Indeed astrocytes and oligodendrocytes are the most abundant cells in WM (based on cell counts), accounting for more than half the volume of a MRI voxel (Walhovd, Johansen-Berg, and Káradóttir 2014). It is thus possible that an increase in size or number of macro-glia cells would have a significant effect on the DW-MRI signal, thus contributing to the positive association here observed between physical lifestyle scores and WM *extra*-neurite fraction (by construction 1 minus intra-neurite density (NODDI f-intra), and specifically, the hindered space outside the neurites prescribed through anisotropic diffusion). Crucially, there exists key histological evidence from animal studies in support of an increase in astrocyte proliferation and in glial fibrillary acidic protein (GFAP) levels (Li et al. 2005; Uda et al. 2006) and in oligodendrocytes number (Luo et al. 2019) in several areas of the rat brain. Together with this previous literature, the findings here reported may suggest a positive relationship between physically active lifestyle and macro-glia cell density across multiple WM tracts, perhaps reflecting a role in providing enhanced metabolic support for neurons. This hypothesis should be tested using imaging alongside more direct measures from ex vivo studies, or using alternative techniques with greater specificity, such as detecting MRS-visible metabolites with greater sensitivity to astrocytes (Brand, Richter-Landsberg, and Leibfritz 1993).

We also report two patterns of RSNs involvement in the mode of brain – physical covariation. We found that a physically active lifestyle was linked with greater connectivity in the parietal cortices and with lower connectivity in the occipital cortices, showing respectively increased and decreased BOLD coupling with all RSNs in more active participants. The same phenotype of a physically active lifestyle was also positively related with greater amplitude in local BOLD fluctuations in the dACC and in the parietal cortices, and with lower amplitude in the occipital cortices. Studying both RSNs amplitude (BOLD variance) and functional connectivity (BOLD covariance) can be important in order to understand possible sources of change and the related neural processes (Duff et al. 2018; Garrett et al. 2010). While greater activity both in functional connectivity and in BOLD amplitude may suggest greater co-activation between the parietal cortices and multiple RSNs across the whole-brain, greater BOLD amplitude with no increase in functional connectivity - as observed in the dACC - may suggest greater local activity that results in a decoupling of the dACC from the rest of brain activity. Greater dACC activity during a cognitive control task was previously associated with higher fitness levels in preadolescent children, with greater dACC activity in the high fit group positively related to accuracy in task performance (Voss et al. 2011). In this study however, we found no significant association between pupil’s scores in the mode of brain – physical covariation and differences in cognitive skills or mental health, thus not allowing us to infer on the cognitive or mental health relevance of this brain pattern.

Overall, our findings lend support to the growing body of evidence demonstrating a close relationship between the body and the brain. Although the relatively small sample size given the number of variables of interest and the possible cluster effect of schools may represent a limitation of this study, here we employed thorough statistical procedures (i.e. block-aware permutation testing and leave-one cluster-out cross-validation) to explicitly deal with this factor, thus producing robust and unbiased statistics. Also, the results here reported are correlational and therefore caution is required in interpreting directionality. Nevertheless our findings suggest that a complex physical phenotype that is influenced by physiology, and lifestyle choices, might have widespread effects on biological processes influencing brain phenotypes. Future studies may test whether improving physical health and fitness through means of activity interventions, promotes diffuse neuroplasticity.

In conclusion, this work provides novel insight into the comprehensive relationship between physically active lifestyle and brain structure and physiology in early adolescence. These findings have broad implications for future research, suggesting novel avenues to study the effect of modifiable lifestyle factors as part of wider brain-body relationships. Understanding how physical pathways may foster healthy human brain development can help us to develop better intervention studies aimed at informing public health and education policies.

## Author contributions

P.S., T.W., and H.J.B. designed the experiment; P.S., T.W., C.W. and N.B. collected and processed the data; M.C., D.P., M.B, S.F. and E.D. contributed to data processing; P.S. analysed the data; T.N., S.S., A.M.W., and G.D. contributed to data analysis; and P.S. and H.J.B. wrote the manuscript. All authors contributed with the interpretation of the findings and in writing the manuscript.

## Acknowledgements

We would like to thank all the Fit to Study investigators (https://www.fit-to-study.org/investigators) for their contributions to the trial. The following individuals helped with data collection: Emma Eldridge, Emily Plester, Emily Curtis, Andy Meaney, Patrick Esser, Johnny Collett, Thomas Smejka, Jack Possee, Oliver Bushnell, Eneid Leika, and Cyrus Goodger. Fit to Study was funded by the Education Endowment Foundation and the Wellcome Trust under their ‘Education and Neuroscience’ Programme (Grant Number 2681). H.J-B is supported by the Wellcome Trust (110027/Z/15/Z) and the Oxford NIHR Biomedical Research Centre. G.D. is supported by the UK MRC (MR/K006673/1). T.E.N. is supported by the Wellcome Trust (100309/Z/12/Z). The Wellcome Centre for Integrative Neuroimaging is supported by core funding from the Wellcome Trust (203139/Z/16/Z). Finally, we would like to thank all of the pupils, and their parents, who took part in, and fully engaged with, each aspect of this brain imaging substudy.

